# The native European *Aedes geniculatus* mosquito species can transmit Chikungunya virus

**DOI:** 10.1101/527382

**Authors:** Jorian Prudhomme, Albin Fontaine, Guillaume Lacour, Jean-Charles Gantier, Laure Diancourt, Enkelejda Velo, Silva Bino, Paul Reiter, Aurélien Mercier

## Abstract

Europe is the world’s leading tourism destination and is receiving every year travelers from areas with active arbovirus transmission. There is thus a threat of mosquito-borne virus emergence in Europe due to the presence of the invasive mosquito vector *Aedes albopictus*. Little attention has been paid about the possible role of indigenous mosquito species as vectors of emerging arboviruses. Here, we assessed the vector competence dynamic of *Ae. geniculatus*, a European anthropophilic mosquito species, for chikungunya virus (CHIKV) in comparison with *Ae. albopictus*.

We revealed that *Ae. geniculatus* was highly susceptible to CHIKV infection and could transmit the virus. By specifically exploring the vector competence dynamic in both mosquito species, we revealed that the cumulative distribution of CHIKV incubation period in *Ae. geniculatus* was delayed by several days as compared to *Ae. albopictus*.

Our results strengthen the importance of considering indigenous species as potential vectors for emerging arboviruses. They also revealed the importance of considering variation in arbovirus dissemination or transmission dynamics in mosquitoes when performing vector competence assays. We will discuss the implications of our results on a CHIKV outbreak dynamic in a theoretical framework.

**Sentence summary:** The European mosquito *Aedes geniculatus* is highly susceptible to CHIKV infection but disseminate and transmit the virus several days later than *Ae. albopictus*.

## Introduction

Chikungunya virus (CHIKV) is a mosquito-borne virus that has emerged from its sylvatic habitat in Africa and is now transmitted in many urban regions world-wide wherever competent vectors, primarily *Aedes aegypti* and *Aedes albopictus*, are present. Three distinct lineages of CHIKV are sporadically causing outbreaks in human population [1, 2].

First detected in Albania and later in Italy, the *Ae. albopictus* species [3] is now established throughout much of the Mediterranean basin and northward as far as Paris [4].

The ever-increasing air travel between Europe and regions with active arbovirus transmission, in combination with travel times that are largely inferior to incubation period, has increased the risk of arboviruses introduction into europe, as illustrated by transmission events of CHIKV in several European countries [5–8]. For example, an outbreak of CHIKV in Italy in 2017 involved more than 420 confirmed cases [5].

*Ae. albopictus* has been implicated as the vector for all European CHIKV outbreaks but an indigenous species, *Aedes geniculatus* (*Olivier, 1791*) is frequently found in ovitraps used for surveillance of *Ae. albopictus* in outbreak areas [9]. Both are tree-hole species that breed in natural containers in woodland as well as in man-made containers in the peri-urban and peri-domestic environment [10, 11]. Their eggs are resistant to desiccation and can overwinter in temperate areas. Both species are day-active, exophilic and feed aggressively on humans and other mammals. The implication of European mosquitoes in the transmission of emerging arboviruses has been poorly investigated despite evidence of transmission by other *Aedes* species outside Europe [2]. We compared the competence of the native European mosquito species *Ae. geniculatus* to transmit CHIKV in comparison with the invasive and reference vector species for CHIKV: *Ae. albopictus*. Importantly, the consideration of the dynamic nature of vector competence revealed that differences of vector competence between mosquito species were due to a time shift in the distribution of extrinsic incubation periods rather than differences in maximum proportion of infectious mosquitoes. The importance of considering indigenous species as potential vectors for arbovirus and the implication, in a theoretical framework, of our results on a CHIKV outbreak dynamic would be discussed.

## Materials and Methods

### Mosquito collection and identification

Mosquitoes from both *Ae. albopictus* and *Ae. geniculatus* species used in this study originated in Tirana, the capital of Albania. Eggs were collected by ovitrap in an urban park (41° 18’36” N; 19° 49’18” E) from July to August 2012. Eggs from both species were hatched at the Institut Pasteur in Paris and reared under standard conditions. Adults from the F0 generation were identified morphologically [12] and 500 individuals of each species wereplaced per cage at 26°C±1°C with 60-70% relative humidity and a light: dark cycle of 16 h: 8 h. Adults were given 10% sucrose solution and females were allowed to engorge with rabbit blood on a membrane feeding apparatus (Hemotek, Discovery Workshops, Lancashire, United Kingdom) to obtain F1 eggs. Batches of eggs were hatched simultaneously to obtain females of the same physiological age for experimental infections. Larvae were reared to the adult stage under the same conditions. All experiments were realized with the F_1_ generation.

### Virus

The isolate CHIKV 0621 was used in this study. This strain is phylogenetically close to an Italian strain with an amino acid change (A226V) in the envelope glycoprotein E1 [13–15] [GenBank accession number DQ443544]. The virus had been passaged three times in C6/36 cells prior to use in the experiments. Virus titration was performed by focus-forming assay (FFA) as previously described [16]. The titer of the frozen virus stock was estimated as 10^9^ focus-forming units per mL (FFU/mL). All infectious experiments were conducted in a BSL-3 insectary (Institut Pasteur, Paris).

### Mosquito infections

Females 9-10 days old were deprived of sucrose solution 24 hours before experimental infection. The infectious blood-meal consisted of 1 mL of viral suspension, 2 mL of washed rabbit erythrocytes supplemented with adenosine triphosphate (10 mM) as a phagostimulant. Blood feeding was by an artificial feeding apparatus (Hemotek) covered with pig intestine. The final virus titer in blood meal was 10^8^ FFU/mL corresponding to the viral load encountered in some patients [17, 18]. Feeders were maintained at 37°C and placed on top of the mesh of a plastic box containing 60 females. After 15 minutes of feeding (to minimize the effect of virus degradation in the infectious blood meal), mosquitoes were cold anesthetized and sorted on ice. Fully engorged females were transferred to cardboard containers and maintained with 10% sucrose in an environmental growth-cabinet set at 28°C ±1°C, 80% humidity, and a 16 h: 8 h light regime.

### Vector competence and virus titration

CHIKV infection, systemic infection and transmission was determined at 3, 5, 7, 10, 12, 14 and 20 days post virus exposure (DPE) for both species. Presence of CHIKV in bodies indicates a midgut infection, while presence of virus in heads and saliva indicates a systemic (disseminated) infection and virus transmission, respectively. Saliva collection was performed using the forced salivation technique [19].

Virus titration was performed by visualizing infectious foci on a sub confluent culture of C6/36 cells by indirect immunofluorescence using 10-fold serial sample dilutions as previously described [20] with a minor modification: After fixation, cells were washed three times in PBS and incubated for 1 □hour at 37□°C with 50 □μL/well of mouse ascetic fluid specific to CHIKV (primary antibody) diluted 1:1000 in PBS□+□1% bovine serum albumin (BSA) (Interchim, Montluçon, France). This mouse ascetic fluid was made by the Centre National de reference des arbovirus of the Institut Pasteur in March 2013 and provided by Pr. Despres Philippe.

### Statistical analysis

The time-dependent effect of the mosquito species on mosquito midgut infection, systemic infection and virus transmission was analysed by Firth’s penalized likelihood logistic regression by considering each phenotype as a binary response variable. Penalized logistic regression, implemented through the *logistf* R package [21] was used to solve problem of separation that can occur in logistic regression when (i) the outcome has high prevalence, (ii) when all observations have the same event status for a combination of predictors or (iii) when a continuous covariate predict the outcome too perfectly. A full-factorial generalized linear model that included the time post virus exposure and the mosquito species was fitted to the data with a binomial error structure and a logit link function. Statistical significance of the effects was assessed by an analysis of deviance.

Virus titres in mosquito’s bodies, heads and saliva were compared between species by a Mann-Whitney-Wilcoxon rank sum test stratified on the time post-exposure as implemented in the *wilcox_test* function from the *coin* R package [22]. The effect of time post exposure on each quantitative phenotype for each species was assessed using a Kruskal-Wallis rank sum test. Time post exposure was converted into ordered factors to implement both tests.

The intra-host dynamic of systemic infection was assessed by a global likelihood function for each mosquito species as described in Fontaine *et al*, 2018 [23]. Probabilities of systemic infection at each time point post virus exposure were estimated with a 3-parameter logistic model. The probability of systemic infection at a given time point (*t*) is governed by *K*: the saturation level (the maximum proportion of mosquitoes with a systemic infection), *B*: the slope factor (the maximum value of the slope during the exponential phase of the cumulative function, scaled by *K*) and *M*: the lag time (the time at which the absolute increase in cumulative proportion is maximal). For easier biological interpretation, *B* was transformed into Δt, which correspond to the time required to rise from 10% to 90% of the saturation level with the formula: Δt = ln (81) / *B*. For each mosquito species, the *subplex* R function [24] was used to provide the best estimates of the three parameters to maximize the global likelihood function (i.e., the sum of binomial probabilities at each time point post virus exposure). This method accounts for differences in sample size when estimating parameters values.

## Results

A total of 150 *Ae. geniculatus* and 141 *Ae. albopictus* engorged females were analysed, considering the mosquito mortality during the experiment. High midgut infection prevalences (>85%) were obtained for both species from the first time point after virus exposure. With 100% midgut infection among the exposed mosquitoes, *Ae. geniculatus* was highly susceptible to CHIKV infection (Supplementary figure 1). Midgut infection prevalences were not influenced by the time post exposure (analysis of deviance, *p*-value = 0.0659) and a significantly higher midgut infection prevalence was observed in *Ae. geniculatus* vs *Ae. albopictus*(analysis of deviance, *p*-value = 0.0438).

CHIKV titres in bodies were significantly higher in *Ae. geniculatus* vs *Ae. albopictus* (stratified Mann-Whitney-Wilcoxon rank sum test, *p*-value < 2.2e-16), with approximately one log 10 difference between values average across time points (5.1 *vs* 4.2 log 10 FFU/mL for *Ae. geniculatus* and *Ae. albopictus*, respectively). Virus titres in bodies were significantly different across time points post virus exposure in both species (Kruskal-Wallis rank sum test, *p*-value = 0.0002402 and 2.984e-15 for *Ae. geniculatus* and *Ae. albopictus*, respectively).

Systemic infection, as measured by the head infection prevalence, was significantly influenced by the time post exposure (analysis of deviance, *p*-value = 5.43e-13) and the mosquito species (analysis of deviance, *p*-value = 4.10e-15) but the time post exposure effect was not significantly different across the mosquito species according to logistic regression (analysis of deviance, interaction term, *p*-value= 0.6). The intra-host dynamic of systemic CHIKV infection was quantified in both mosquito species by fitting a 3 parameters logistic model that assumes a sigmoidal distribution of the cumulative proportion of mosquitoes with a systemic infection over time. Both species presented a saturation level (K, or maximum proportion of infected mosquitoes with a systemic infection) equal to 100% (Figure 1). However, this saturation level was later for *Ae. geniculatus* than for *Ae. albopictus*. Indeed, an estimated 14.6 days were needed for *Ae. geniculatus* to go from 10% to 90% of the saturation level whereas 100% of the saturation level was reached in less than one day for *Ae. albopictus*. The lag time *M*, which can represent a proxy for the extrinsic incubation period-50 (EIP-50—the time required for the systemic infection prevalence to reach 50% of the saturation level)—was 2.7 days in *Ae. albopictus* mosquitoes compared to 7.6 days in *Ae. geniculatus* (Figure 1).

**Figure 1:**
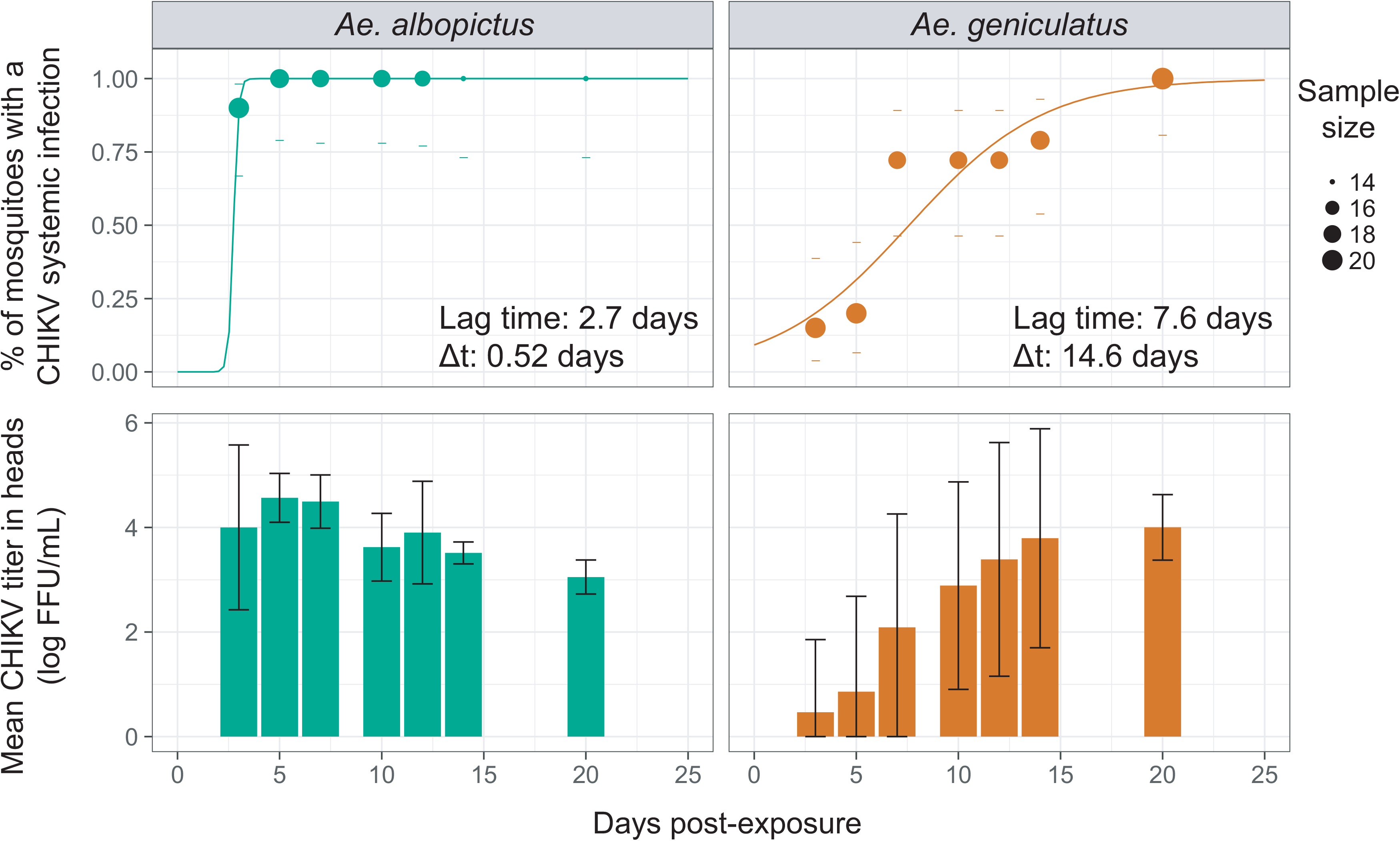
Systemic infection kinetic for *Ae. albopictus* and *Ae. geniculatus* species infected with CHIKV. The upper panel shows the cumulative prevalence of systemic infection over time post CHIKV exposure. Data points represent the observed prevalence at each time point with their size being proportional to the sample size. Dashes represent the 95% confidence interval of the prevalence. The fitted values obtained with a 3-parameter logistic model are represented for each species by a coloured line. The lag time and rising time estimates are represented for each species. The lower panel shows CHIKV titres in mosquito heads measured at each time points after oral exposure to the virus.

CHIKV titres in heads were significantly lower in *Ae. geniculatus* vs *Ae. albopictus* (stratified Mann-Whitney-Wilcoxon rank sum test, *p*-value = 3.3e-4), with approximately 1.3 log _10_ difference between values average across time points (2.5 *vs* 3.8 log 10 FFU/mL for *Ae. geniculatus* and *Ae. albopictus*, respectively). Virus titres in heads were also significantly different across time points post virus exposure in both species (Kruskal-Wallis rank sum test, *p*-value = 2.621e-07 and 1.781e-10 for *Ae. geniculatus* and *Ae. albopictus*, respectively).

Virus transmission prevalences were measured by assessing the presence of infectious virus in mosquito saliva. Virus transmission prevalences were significantly different between mosquito species (analysis of deviance, *p*-value = 0.0038) and the effect of time post exposure varied significantly between mosquito species (analysis of deviance, interaction term, *p*-value = 3.05e-4). It was not possible to model the cumulative virus transmission prevalence over time for the *Ae. albopictus* species because of a lack of saturation level. Proportion of virus in saliva clearly collapsed from day 5 post virus exposure. The same phenomenon could be observed for *Ae. geniculatus*. However, the 3-parameter model could nonetheless fit the data, giving the following estimates: 70% of saturation level, an EIP-50 of 8 days and a rising time of 6.2 days (Supplementary figure 2).

CHIKV titres in saliva were significantly higher in *Ae. geniculatus* vs *Ae. albopictus* (stratified Mann-Whitney-Wilcoxon rank sum test, *p*-value = 1.8e-3), with 0.24 log 10 difference between values average across time points (0.51 *vs* 0.27 log 10 FFU/mL for *Ae. geniculatus* and *Ae. albopictus*, respectively). Titres in heads were significantly different across time points after virus exposure in mosquitoes from the *Ae. geniculatus* species only (Kruskal-Wallis rank sum test, *p*-value = 4e-03 and 0.14 for *Ae. geniculatus* and *Ae. albopictus*, respectively). Virus titres in saliva were clearly increasing over time in mosquitoes from the *Ae. geniculatus* species.

## Discussion

Chikungunya virus (CHIK) is one of four arboviruses withdengue virus (DENV), Zika virus (ZIKV) and yellow fever virus (YFV) that can be sustained in a human-vector-human transmission cycle. All four originated in wild primates but can cycle in the urban environment, transmitted by peridomestic mosquitoes. Intercontinental travel and trade were historically involved in the transmission of these viruses world-wide, sometimes involving recruitment of new local vectors [25, 26]. Thus, the YFV, originating in Africa, was introduced to the Americas during the slave trade,where it entered new transmission cycles involving autochthonous arboreal species, such as *Haemagogus janthinomys* and mosquito from the *Sabethes* genus, and non-human primates in primary tropical rain forest. The virus then regularly propagated from the jungle into urban transmission cycles [27].

Europe is the world’s leading tourism destination with 671 million arrivals (50% of all international tourists) in 2017, a number that continues to increase. [28]. Travellers with arbovirus infections [29–35], including CHIKV [36] have initiated autochthonous transmission [5, 6] vectored by the invasive mosquito *Ae. albopictus*. At the same time, Europe harbours native mosquito species potentially able to transmit emerging arboviruses. Yet, only two studies have assessed the potential for CHIKV infection in six indigenous species from the south of France and Italy (*Culex pipiens, Aedes caspius, Aedes detritus, Aedes vexans, Aedes vittatus*, and *Anopheles maculipennis*). The four *Aedes* species were susceptible to virus infection, but the transmission potential of infectious viral particles in the mosquito saliva was not investigated. These insects represent a small portion of all European anthropophilic mosquitoes.

Our study has revealed the ability of the European mosquito *Ae. geniculatus* to transmit CHIKV experimentally. *Aedes geniculatus* was revealed to be highly susceptible to CHIKV infection. Midgut infection by an arbovirus is achieved very rapidly after exposure to the infectious blood meal when infectious virus particles are still in contact with the midgut wall cells. Virus dissemination in secondary tissues can only occur in mosquitoes with infected midgut and is, in contrast to the infection phenotype, a dynamic process scaled in a day unit that can have epidemiological significance. Variation in DENV dissemination dynamics in mosquitoes was reported to significantly affects the risk and magnitude of dengue outbreaks [23]. We revealed that both species were completely able to disseminate CHIKV in their secondary tissues. However, this 100% saturation level of infected mosquitoes with a disseminated infection was reached in *Ae. albopictus* 15 days earlier than *Ae. geniculatus*. The estimated virus dissemination lag time, that can be considered as a proxy for the EIP-50 (extrinsic incubation period in days until 50% of maximum infectiousness), was nearly 8 days for *Ae. geniculatus* whereas is was less than 3 days for *Ae. albopictus*. Combined with vector longevity, EIP is the most powerful contributor to vectorial capacity according to the Ross-MacDonald equation [37]. Vectorial capacity defines the number of secondary infections expected to occur from the introduction of a single infection in a naive population [38]. The longer it would take for a virus to disseminate from the infected midgut to the saliva, the fewer opportunities the mosquito would have to transmit the virus to a human host before its death. This theory can be thwarted if *Ae. geniculatus* mosquito’s life span is equal or longer than *Ae. albopictus* one [39]. *Ae. geniculatus* might then be involved in autochthonous CHIKV transmissions in Europe in complementation with or in absence of *Ae. albopictus*. In addition, possibility of rapid virus adaptation to new vectors was already observed [40, 41]. CHIKV evolution toward restricted EIP duration in this autochthonous species would greatly increase the epidemic potential of this vector-virus couple.

The biology of *Ae. geniculatus* is poorly studied [10]. From what is known, the species can bite aggressively both humans and animals mainly during daylight hours [10, 57] (and personal observation). Its larvae are commonly found in mature trees’ holes [42]. Adults often coexist with *Ae. albopictus* in Europe but population densities of *Ae. geniculatus* generally didn’t reached those of *Ae. albopictus* in peri-urban areas [64]. More data are needed to fully characterise the level of contact with human hosts. As an anecdote, we captured *Ae. geniculatus* on Jim Morrison grave, one of the most visited graves by tourists of all nationalities in Pere Lachaise cemetery, in the center of Paris, where millions of people come every year, strengthening the threat for this species in local virus transmission. Its flight dispersion is not considered superior than *Ae. albopictus*. Consequently, this species would not carry viruses outside of the area treated in the frame of the vector control intervention [59]. Arbovirus vertical transmission is a rare event, and its occurrence hazard is correlated with vector density and the surveillance of virus. If higher CHIKV vertical transmission rates are achieved in *Ae. geniculatus* species, vertical transmission of CHIKV in overwintering *Aedes* mosquitoes might contribute to the maintenance of this virus during winter. Populations of *Ae. geniculatus* are generally monovoltine (ie. one generation per year), and their diapause termination is asynchronous [10], enabling a reintroduction of the virus later in the following year when the population of the primary vector *Ae. albopictus* is already high. It seems therefore necessary to improve the knowledge on this species, including longevity, flight and host-seeking behaviour, diapause and virus overwintering to fully assess its epidemic potential. At last, temperature is strongly influencing vector competence. It can be of interest to compare vector competence for CHIKV between both mosquito species at lower temperatures.

Our study has several limitations. First, our estimates of virus dissemination dynamic can be biased. They rely on the modelisation of the measured cumulative proportion of mosquitoes with a disseminated infection over time. Concerning *Ae. geniculatus*, the modelized proportion is not null at the time of mosquito exposure to the virus as it should be. The number and repartition of time points and the proportions accuracy (that depend on sample size at each time point) are influential in determining accurate parameter estimates of the virus dissemination dynamic. This method can nonetheless provide realistic estimates and offer the possibility to consider the dynamic nature of vector competence. Importantly, this allows to disentangle the incubation period effect from the maximum proportion of systemic virus infection. Then, the transmission dynamic estimates must be considered with caution because the method to detect viruses in mosquito saliva do not distinguish true negatives from negatives resulting from mosquito that did not expectorate saliva. This can explain decrease of transmission rates for *Ae. albopictus*. Alternatively, this decrease could be explained by the fact that infected mosquitoes died before non-infected mosquitoes or that oldest mosquitoes begin to clear the infection through immune function [43] or by the natural death of virions. It can however be concluded that both species can transmit infectious CHIKV particles as soon as 3 and 7 days post exposure for *Ae. albopictus* and *Ae. geniculatus*, respectively.

CHIKV is in expansion throughout the world and Europe is not spared. Since the CHIKV outbreak in Northern Italy in 2007, and more recently the isolated cases of chikungunya and dengue recorded in France and Croatia from 2010 to 2013 [30, 44], it has been acknowledged that Europe is vulnerable to local transmission of “tropical” arboviruses. Epidemics risk is in connection with the steady increase of imported cases of Aedes-bornes viruses as well as with the European expansion of *Ae. albopictus* [45]. So far, studies on the epidemiology of the arboviral diseases chikungunya and dengue in European countries have focused on the invasive “Asian tiger mosquito” without considering the potential role for indigenous vector species. These results show the importance of considering European indigenous species to assess overall risk of arbovirus transmission in Europe.

Assessing the vector competence of the different European mosquito species, but also other regions around the world potentially exposed to arboviral outbreak, for other arboviruses or other strains, will help to anticipate patterns of transmission of arbovirus as CHIKV and DENV and the relative contribution of different vector species to virus’s amplification and persistence (transmission and reservoir) in these areas. Moreover, we recommend to monitor on the field all mosquito species, whether native or invasive, during surveillance programs on future outbreaks, to further characterize the autochthonous mosquito fauna that can be potentially involved in arbovirus transmission.

## Acknowledgments

We thank Pr. Despres Philippe for providing technical assistance. This study was funded by EU grant FP7-261504 EDENext (http://www.edenext.eu). The contents of this publication are the sole responsibility of the authors and do not necessarily reflect the views of the European Commission.

## Tables

**Table 1:** Body infection rate (BIR), systemic infection rate (SIR) and transmission rate (TR) at different days post-infection of *Ae. geniculatus* and *Ae. albopictus* exposed to CHIKV. Mosquitoes were contained in cardboard boxes after virus exposure. For all surviving mosquitoes per box and time point, we measured three parameters describing vector competence: (i) mosquito infection as measured by the presence of viral infectious particles in the body (abdomen and thorax): BIR, (ii) virus dissemination as measured by the presence of viral infectious particles in mosquito head: SIR, and (iii) transmission efficiency as measured by the number of mosquitoes with viral infectious particles in their saliva: TR.

| Day pi | *Ae. geniculatus* |  |  | *Ae. albopictus* |  |  |
| --- | --- | --- | --- | --- | --- | --- |
|  | BIR<sup>a</sup> | SIR<sup>b</sup> | TR<sup>c</sup> | BIR<sup>a</sup> | SIR<sup>b</sup> | TR<sup>c</sup> |
| 3 | 100 (20) | 15 (20) | 0 (3) | 100 (20) | 90 (20) | 27.8 (18) |
| 5 | 100 (20) | 20 (20) | 0 (4) | 95 (20) | 100 (19) | 63.1 (19) |
| 7 | 100 (18) | 72.2 (18) | 30.8 (13) | 100 (18) | 100 (18) | 38.9 (18) |
| 10 | 100 (18) | 72.2 (18) | 38.5 (13) | 100 (18) | 100 (18) | 27.8 (18) |
| 12 | 100 (18) | 72.2 (18) | 84.6 (13) | 100 (17) | 100 (17) | 23.5 (17) |
| 14 | 100 (19) | 78.9 (19) | 73.3 (15) | 93.3 (15) | 100 (14) | 28.6 (14) |
| 20 | 100 (21) | 100 (21) | 61.9 (21) | 87.5 (16) | 100 (14) | 14.3 (14) |

<sup>a</sup> Percent of mosquitoes with infected body among all exposed mosquitoes.
<sup>b</sup> Percent of mosquitoes with infected head among body infected mosquitoes.
<sup>c</sup> Percent of mosquitoes with infectious saliva among mosquitoes with a systemic infection (mosquitoes that disseminate the virus beyond the midgut barrier).
The number of mosquitoes analysed is displayed in brackets.

---

**Supplementary figure 1:**
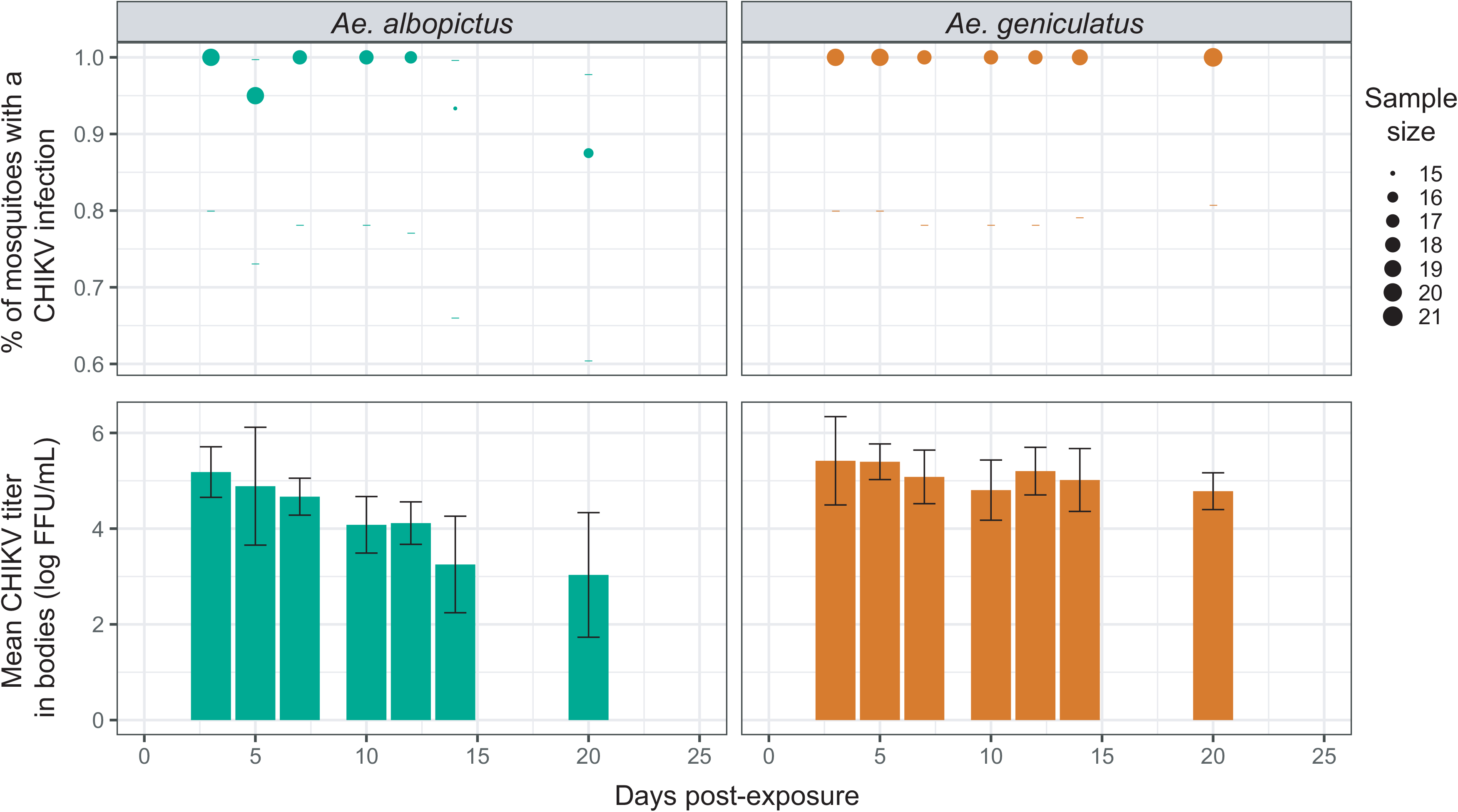
Midgut infection for *Ae. albopictus* and *Ae. geniculatus* species infected with CHIKV. The upper panel shows the percentage of midgut infection over time post CHIKV exposure. Data points represent the observed prevalence at each time point with their size being proportional to the sample size. Dashes represent the 95% confidence interval of the prevalence. The lower panel shows CHIKV titres in mosquito bodies measured at each time points after oral exposure to the virus.

**Supplementary figure 2:**
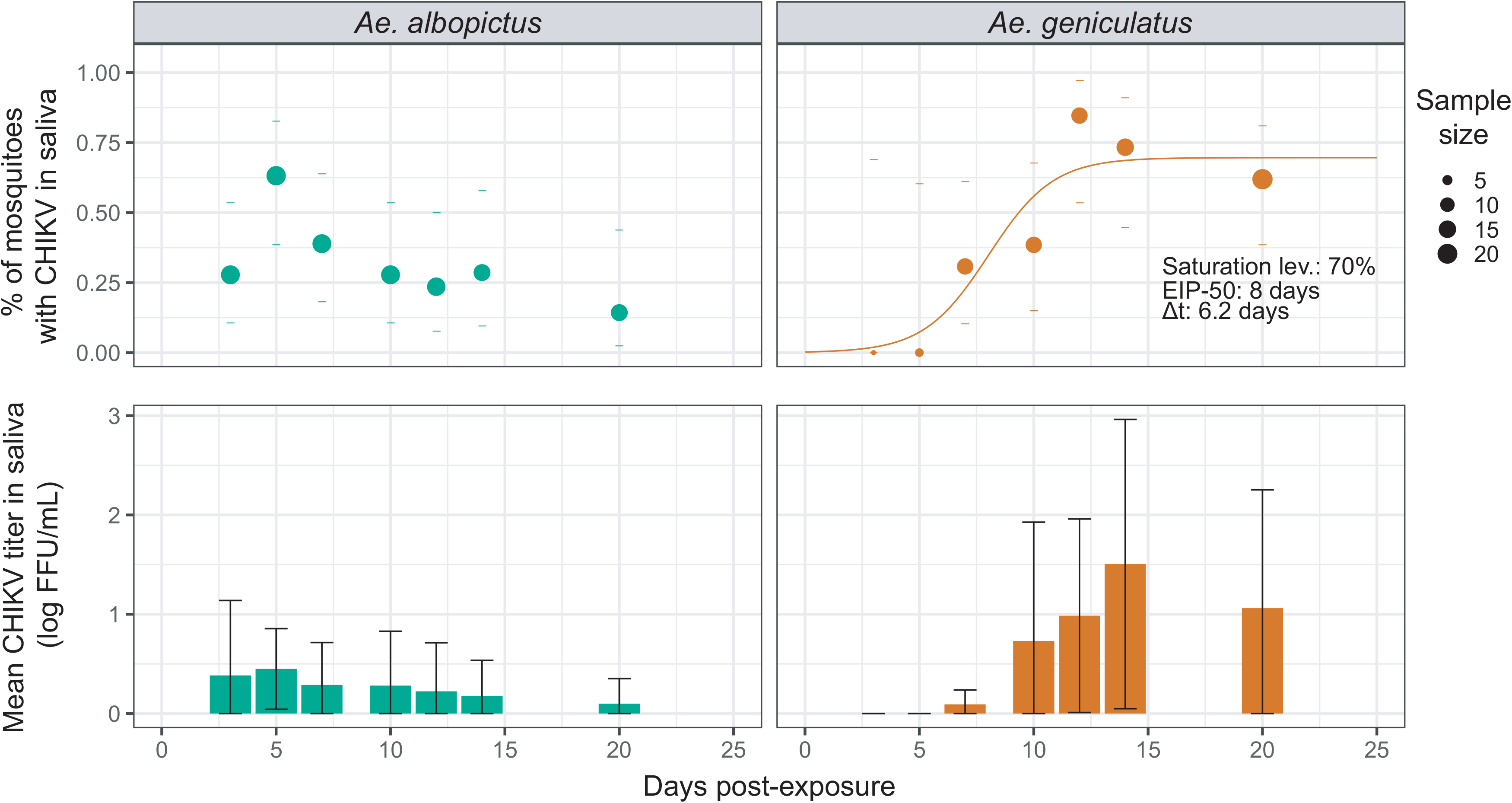
Saliva infection for *Ae. albopictus* and *Ae. geniculatus* species exposed to CHIKV. The upper panel shows the percentage of saliva infection over time post CHIKV exposure. Data points represent the observed prevalence at each time point with their size being proportional to the sample size. Dashes represent the 95% confidence interval of the prevalence. EIP-50 and rising time estimates are represented for *Ae. geniculatus* species. The lower panel shows CHIKV titres in mosquito saliva measured at each time points after oral exposure to the virus.

